# Development of a Triazolobenzodiazepine-Based PET Probe for Subtype-Selective Vasopressin 1A Receptor Imaging

**DOI:** 10.1101/2021.02.09.430516

**Authors:** Ahmed Haider, Zhiwei Xiao, Xiaotian Xia, Jiahui Chen, Richard S. Van, Shi Kuang, Chunyu Zhao, Jian Rong, Tuo Shao, Perla Ramesh, Appu Aravind, Yihan Shao, Chongzhao Ran, Larry J. Young, Steven H. Liang

## Abstract

**Objectives:** To enable non-invasive real-time quantification of vasopressin 1A (V1A) receptors in peripheral organs, we sought to develop a suitable PET probe that would allow specific and selective V1A receptor imaging *in vitro* and *in vivo*.

**Methods:** We synthesized a high-affinity and -selectivity ligand, designated compound **17**. The target structure was labeled with carbon-11 and tested for its utility as a V1A-targeted PET tracer by cell uptake studies, autoradiography, in vivo PET imaging and ex vivo biodistribution experiments.

**Results:** Compound **17** (PF-184563) and the respective precursor for radiolabeling were synthesized in an overall yield of 49% (over 7 steps) and 40% (over 8 steps), respectively. An inhibitory constant of 0.9 nM towards the V1A receptor was measured, while excellent selectivity over the related V1B, V2 and OT receptor (IC_50_ >10,000 nM) were obtained. Cell uptake studies revealed considerable V1A binding, which was significantly reduced in the presence of V1A antagonists. Conversely, there was no significant blockade in the presence of V1B and V2 antagonists. *In vitro* autoradiography and PET imaging studies in rodents indicated specific tracer binding mainly in the liver. Further, the pancreas, spleen and the heart exhibited specific binding of [^11^C]**17** ([^11^C]PF-184563) by *ex vivo* biodistribution experiments.

**Conclusion:** We have developed the first V1A-targeted PET ligand that is suitable for subtype-selective receptor imaging in peripheral organs including the liver, heart, pancreas and spleen. Our findings suggest that [^11^C]PF-184563 can be a valuable tool to study the role of V1A receptors in liver diseases, as well as in cardiovascular pathologies.

## Introduction

Vasopressin 1A (V1A) receptors are transmembrane proteins that belong to the superfamily of G-protein-coupled receptors (GPCRs). [1] Notably, V1A receptor activation is triggered by an endogenous nonapeptide, arginine vasopressin (AVP), which is released from the posterior pituitary gland into the blood stream. [2, 3] Previous studies revealed that V1A receptors are expressed in the liver, cardiovascular system, kidney, pancreas, spleen, adipose tissue and the central nervous system (CNS). [4-6] Given the emerging focus on the role of V1A receptors in neurological pathologies, a plethora of studies outlined their distinct distribution in brain sections of different species.[7-9] Despite 83% of structural homology between rat and human V1A receptors, studies have unveiled substantial species-differences in the *N-*terminal domain that is involved in ligand binding. [10, 11] To this end, significant species-dependent variability has been observed for the binding affinities of several synthetic V1A ligands.

V1A receptors have initially been linked to autism spectrum disorder (ASD) [12, 13], Huntington’s disease [14, 15], primary aldosteronism [16] and dysmenorrhea [17]. As such, strenuous drug discovery efforts led to the discovery of several potent antagonists, of which some have entered the clinical arena (**Figure 1**). YM471 was identified as a potent and long-acting V1A antagonist in rats, rendering it a useful tool to further elucidate the physiological and pathophysiological roles of V1A receptors. [18] Later, Guillon et. al reported on SRX246 and SRX251 as novel orally active V1A antagonists with subnanomolar affinities and CNS activity. As such, oral dosing of SRX246 and SRX251 led to brain levels that were ∼100-fold higher than the respective inhibitory constants (K_i_ values). [19] Notwithstanding the potent anxiolytic effects of another ligand, JNJ-17308616, in preclinical experiments, as well as the high binding affinities towards both human and rat V1A receptors, no clinical studies with JNJ-17308616 have been reported to date. [20] Another potent V1A antagonist, YM218, exhibited a Ki value of 0.18 nM and has proven to attenuate vasopressin-induced growth responses of human mesangial cells. Accordingly, YM218 was suggested as a potential tool for investigating the role of V1A in the development of renal disease.[21, 22] As for the clinically tested candidates, relcovaptan (SR49059) [23] showed favorable effects in patients with dysmenorrhea [24] and primary aldosteronism. [16, 25] Further, SRX246 was tested for the treatment of Huntington disease in a clinical phase II study. [14, 15] A CNS-targeted V1A antagonist, RG7713 (RO5028442) [26], exhibited beneficial effects in the treatment of abnormal affective speech recognition, olfactory identification and social communication in adult ASD patients. [27] Further, the triazolobenzodiazepine, balovaptan (RG7314) showed encouraging results for the treatment of ASD in a clinical phase II study, which led to a Breakthrough Therapy Designation by the U.S. FDA in January 2018. [28] Despite these encouraging findings, the potential of V1A antagonists for the treatment of liver and cardiovascular is largely unexplored. This has, at least in part, been attributed to the lack of appropriate tools for *in vivo* imaging of V1A receptors in peripheral organs.

**Figure 1.**
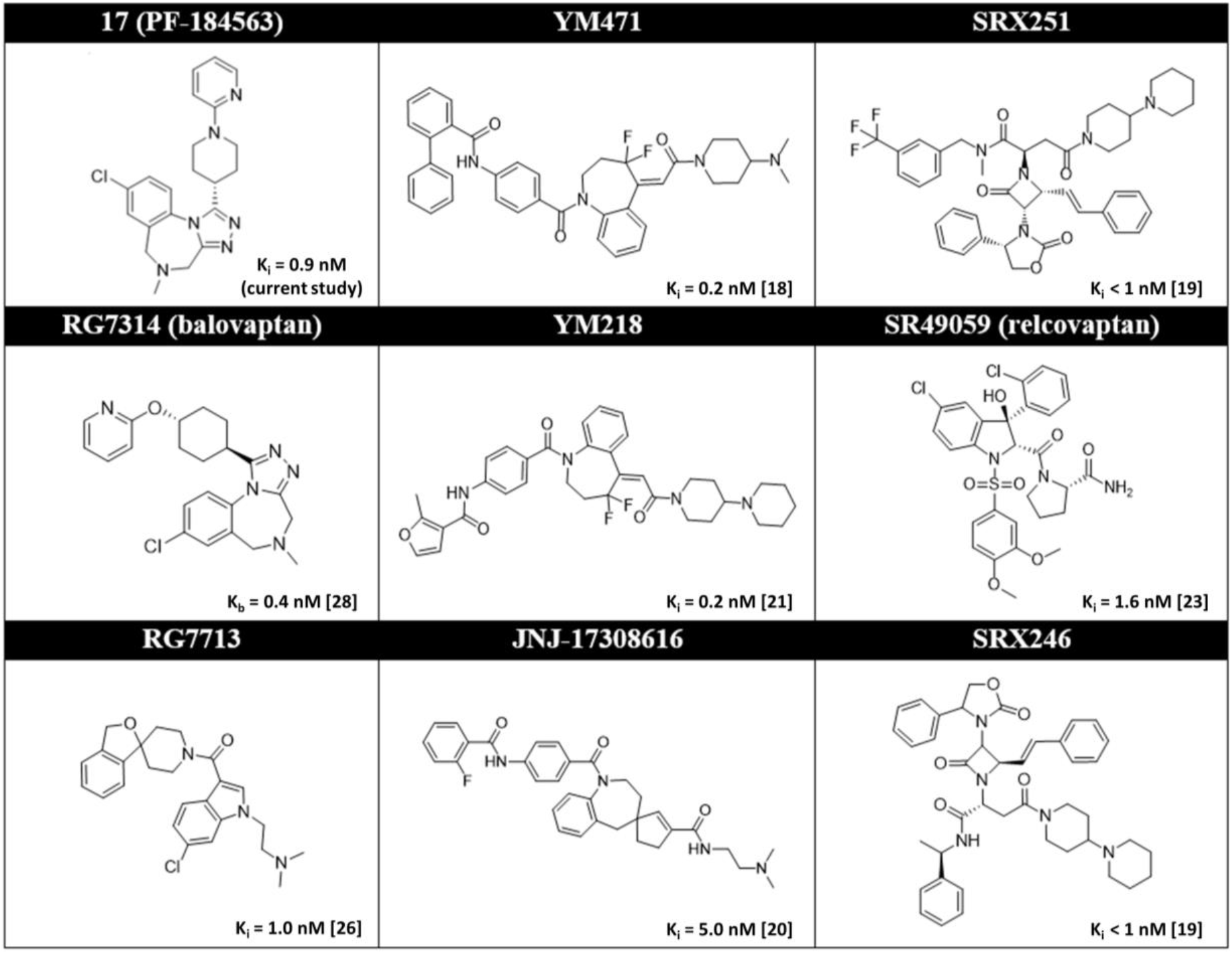
Selected Vasopressin 1A (V1A) receptor antagonists.

Positron emission tomography (PET) is a powerful non-invasive imaging modality that allows real-time quantification of biochemical processes. Accordingly, PET has been established as a reliable tool for receptor quantification in preclinical and clinical research [29]. Further, it has proven particularly useful in facilitating drug discovery via target engagement studies [30, 31]. *In vivo* imaging and quantification of V1A receptors in the CNS as well as in peripheral organs is crucial to advance our understanding on the versatility of V1A receptor-related pathologies. Nonetheless, the development of a suitable PET radioligand remains challenging, and a clinically validated probe is currently lacking. To address this unmet medical need, radiolabeled analogues of SRX246 were synthesized, however, in vivo studies with these probes have not been reported, potentially owing to the relatively high molecular weight (700-800 Dalton) of this class of compounds – a feature that is known to hamper tracer pharmacokinetics. [32] Naik et al. recently reported on the development of [^11^CH_3_]*(1S,5R)-***1** ((4-(1Hindol-3-yl)-3-methoxyphenyl)((1S,5R)-1,3,3-trimethyl-6-azabicyclo[3.2.1]octan-6-yl)methanone), a V1A-targeted PET ligand with subnanomolar biding affinity that was evaluated in rodents. [33] Despite the overall low brain uptake, specific in vivo binding was observed in the lateral septum of CD-1 mice. Further studies are warranted to assess whether sufficient brain exposure can be achieved with [^11^CH_3_]*(1S,5R)-***1** to detect alterations in V1A receptor density in the CNS. While previous PET studies have exclusively focused on imaging V1A receptors in the CNS, there is a rapidly growing body of evidence suggesting that V1A receptors are implicated in serious peripheral pathologies including liver cirrhosis and portal hypertension [34], heart failure [35], diabetes [36], Raynaud’s disease [35], gastric ulcers [37] and renal disease [21, 22]. Accordingly, we sought to develop a suitable V1A-targeted PET probe that would allow quantitative *in vivo* visualization of V1A receptors in peripheral organs such as the liver, heart, kidney, pancreas and the spleen. Along this line, the previously reported triazolobenzodiazepine, PF-184563, was selected for radiolabeling and biological evaluation based on the subnanomolar binding affinity and the appropriate lipophilicity [38]. In the present study, we performed pharmacological profiling of PF-184563, conducted molecular docking studies, and evaluated the radiolabeled analog, [^11^C]PF-184563 ([^11^C]**17**), by cell uptake studies, in vitro autoradiography, *ex vivo* biodistribution experiments and PET imaging in rodents.

## Results and Discussion

### Chemical synthesis

Target compound **17** (PF-184563) and the respective desmethyl precursor **19** were synthesized as previously reported [38], however, with notable differences as outlined in **Schemes 1** and **2**. Indeed, the adapted synthetic strategy provided a common route for key intermediates **11** and **12** (**Scheme 1**), that were used for the synthesis of reference compound **17** and desmethyl precursor **19**, respectively, as depicted in **Scheme 2**. Further, this approach led to a ∼3-fold increase of the overall yield for the synthesis of PF-184563, as compared to the previous report [38]. Briefly, commercially available benzyl alcohol **1** was chlorinated using sulfuryl chloride and the resulting intermediate **2** was employed in an N-alkylation reaction that afforded amines **3** and **4** in 90-99% yield. Subsequently, N-alkylation of **3** with ethyl bromoacetate in the presence of DIPEA yielded **5** in 93%, while amine protection of **4** with Boc anhydride afforded ester **6** in 78% yield. Reduction of the nitro functionality was followed by an intramolecular amidation, which ultimately led to the formation of the benzodiazepine core structures in **9** and **10**, respectively. Key intermediates **11** and **12** were produced via thiation of the carbonyl group using the Lawesson’s reagent.

**Scheme 1.**
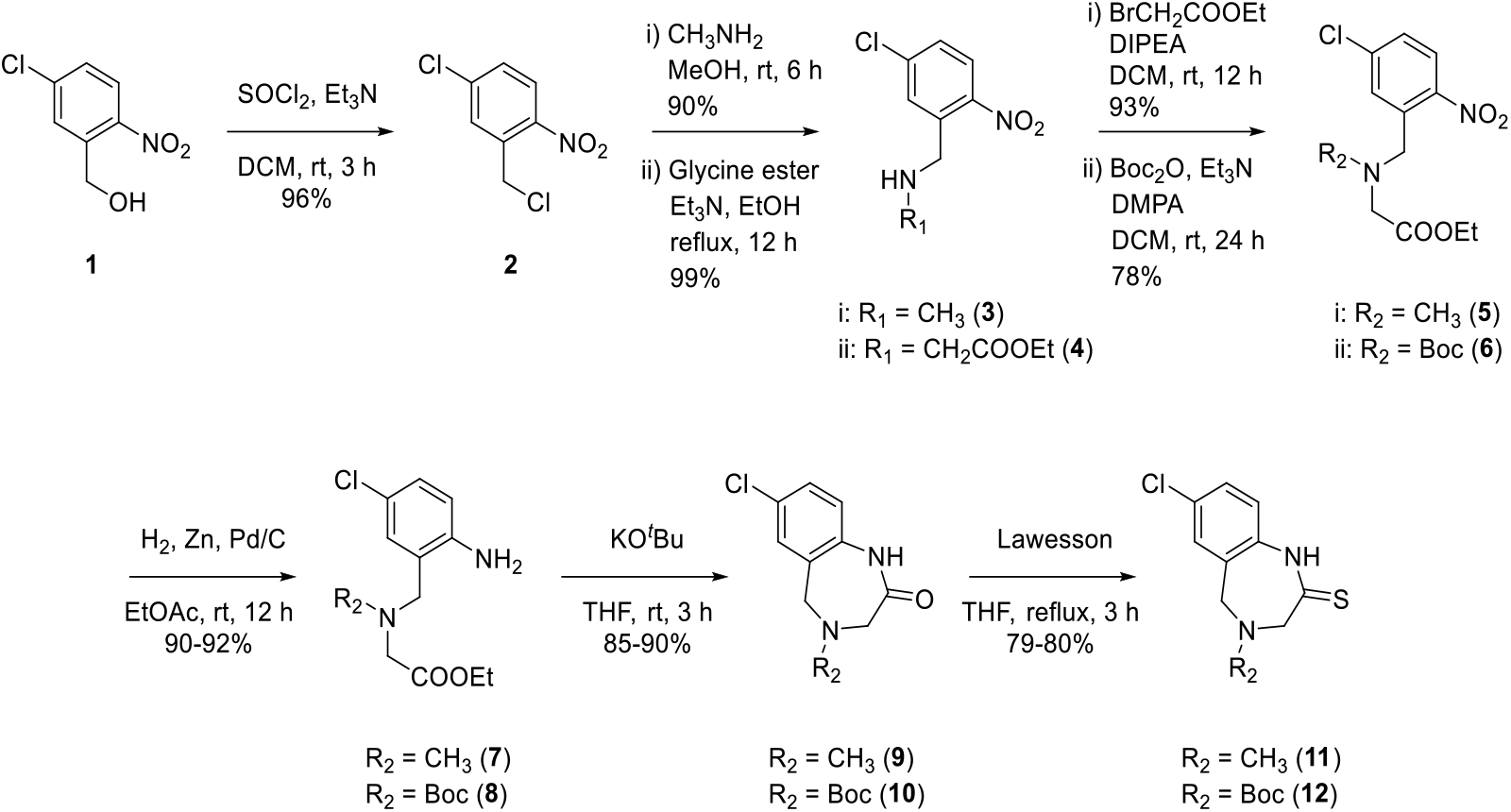
Synthesis of key intermediates **11** and **12**.

Starting from commercially available 3-chloropyridine and ethyl piperidine-4-carboxylate, hydrazine **16** (**Scheme 2**) was synthesized in 83% yield over two steps. The 1,2,4-triazole analogues, reference compound **17** (PF-184563) and intermediate **18**, were prepared by intramolecular cyclization in refluxing n-butanol. Deprotection of **18** under acidic condition afforded precursor **19** in 81% yield. Overall, PF-184563 and the respective precursor **19** were synthesized in a yield of 49% (over 7 steps) and 40% (over 8 steps), respectively.

**Scheme 2.**
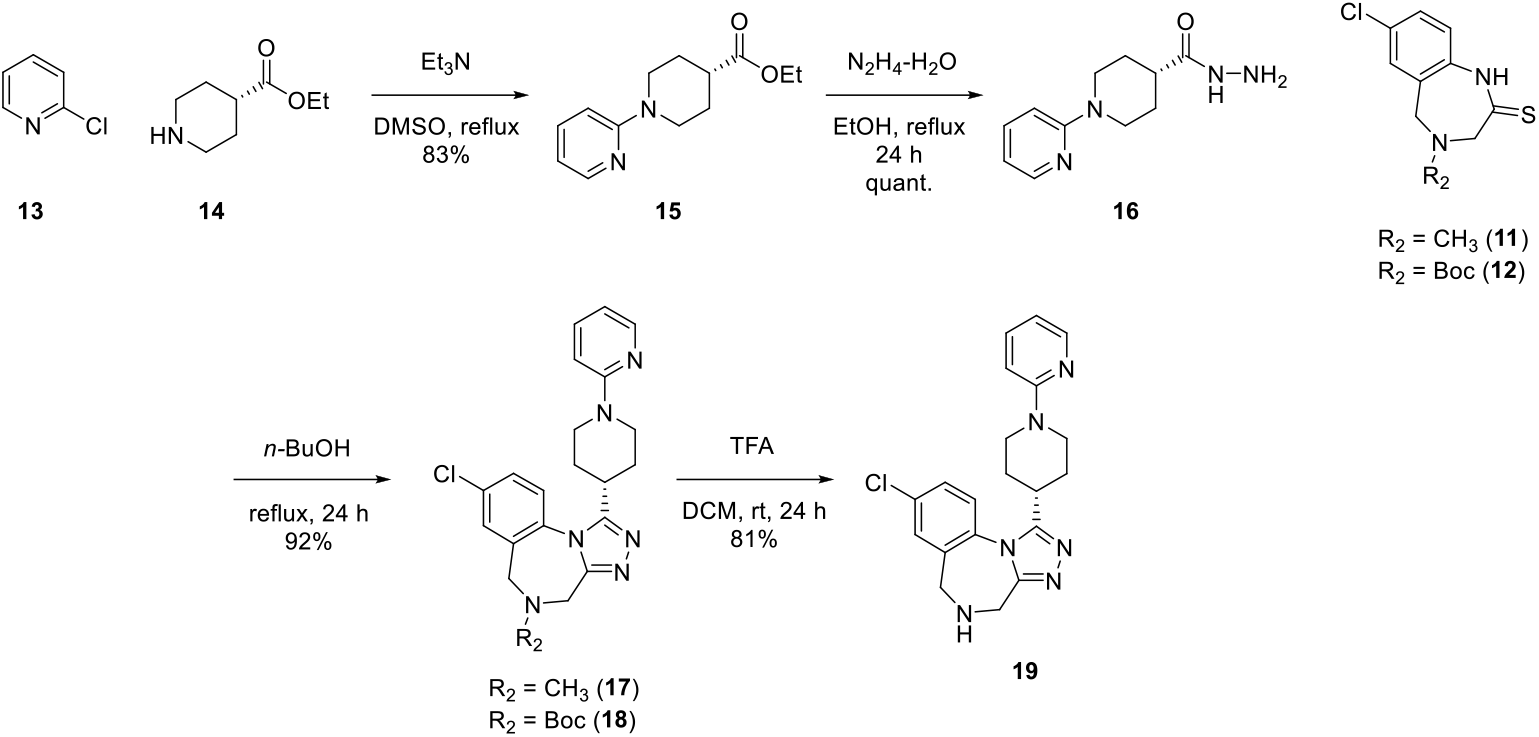
Synthesis of target compound **17** (PF-184563) and precursor **19** for carbon-11 labeling.

### Pharmacology Profiling

Pharmacological properties of target compound **17** were assessed for the human V1A, V1B and V2 and OT, as well as for the hERG potassium channel, the L-type calcium channel and the sodium channel (**Figure 2**). In accordance with previous reports, **17** exhibited a high binding affinity (IC_50_ = 4.7 nM and K_i_ = 0.9 nM) towards the V1A receptor. Indeed, the obtained IC_50_ of 4.7 nM for hV1A was comparable to the previously reported value by Johnson et al. (hV1A IC50: 6.7 nM). [38] Further, the binding affinities for related receptors, including V1B, V2 and oxytocin (OT) receptors, was >10 µM, suggesting a high compound selectivity. Similarly, **17** was deemed inactive against the hERG potassium channel, the L-type calcium channel and the sodium channel in concentrations up to 10 µM. These results confirmed that **17** exhibits high V1A affinity and selectivity.

**Figure 2.**
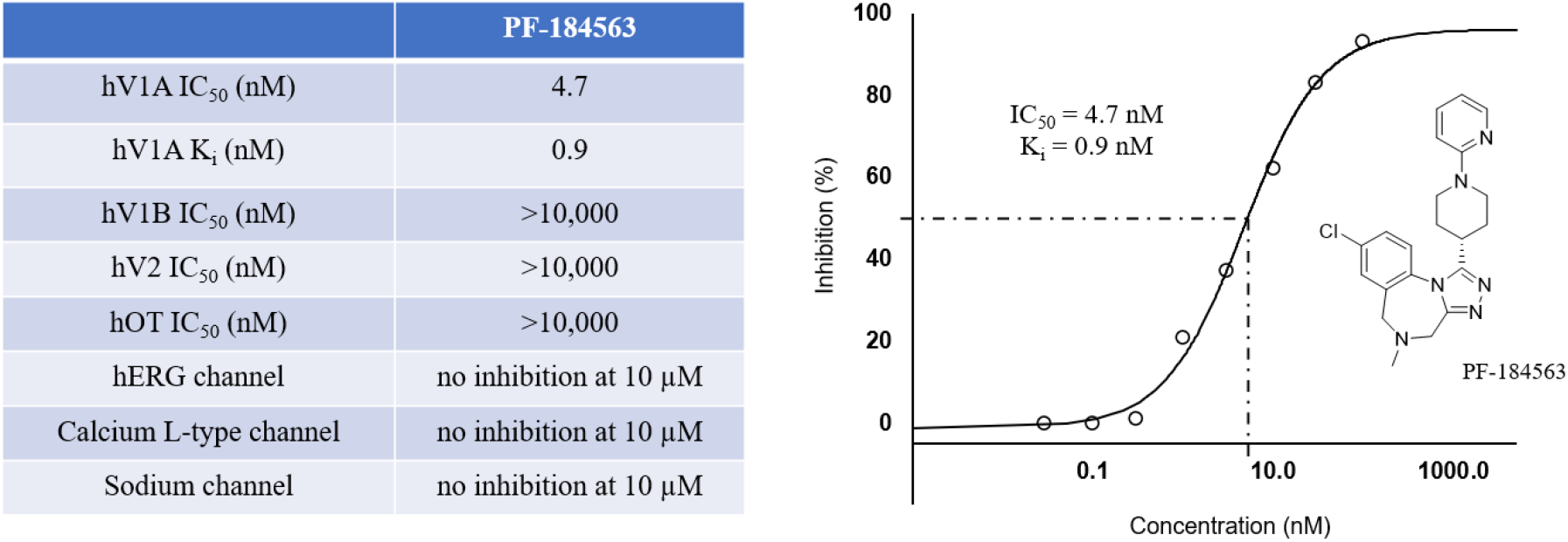
Summary of pharmacological properties for target compound **17** (PF-184563).

### Molecular Docking

To rationalize the subnanomolar binding affinity, molecular interactions of PF-184563 with the V1A receptor were modeled using in silico simulations, as previously reported [39-41]. Since no crystal/NMR structures are available for the V1A receptor, an active state V1A homology model on gpcrdb.org was adopted and subsequently optimized via restrained energy minimization. Various compounds were docked to the V1A binding pocket of the homology model, which is composed of hydrophobic residues. In the docking pose shown in **Figure 3**, the chlorine atom of PF-184563 was found to be in close proximity to the carbonyl groups of T333 (3.115 Å) and C303 (3.025 Å). Similarly, the triazole moiety was near Q131, with the closest distance between the amide hydrogen of Q131 and triazole nitrogens being 2.76 Å. These findings support the concept that hydrogen bond interactions between Q131 and the triazole moiety of PF-184563 may stabilize the binding pose, thereby contributing to the low nanomolar affinity observed towards the V1A receptor.

**Figure 3.**
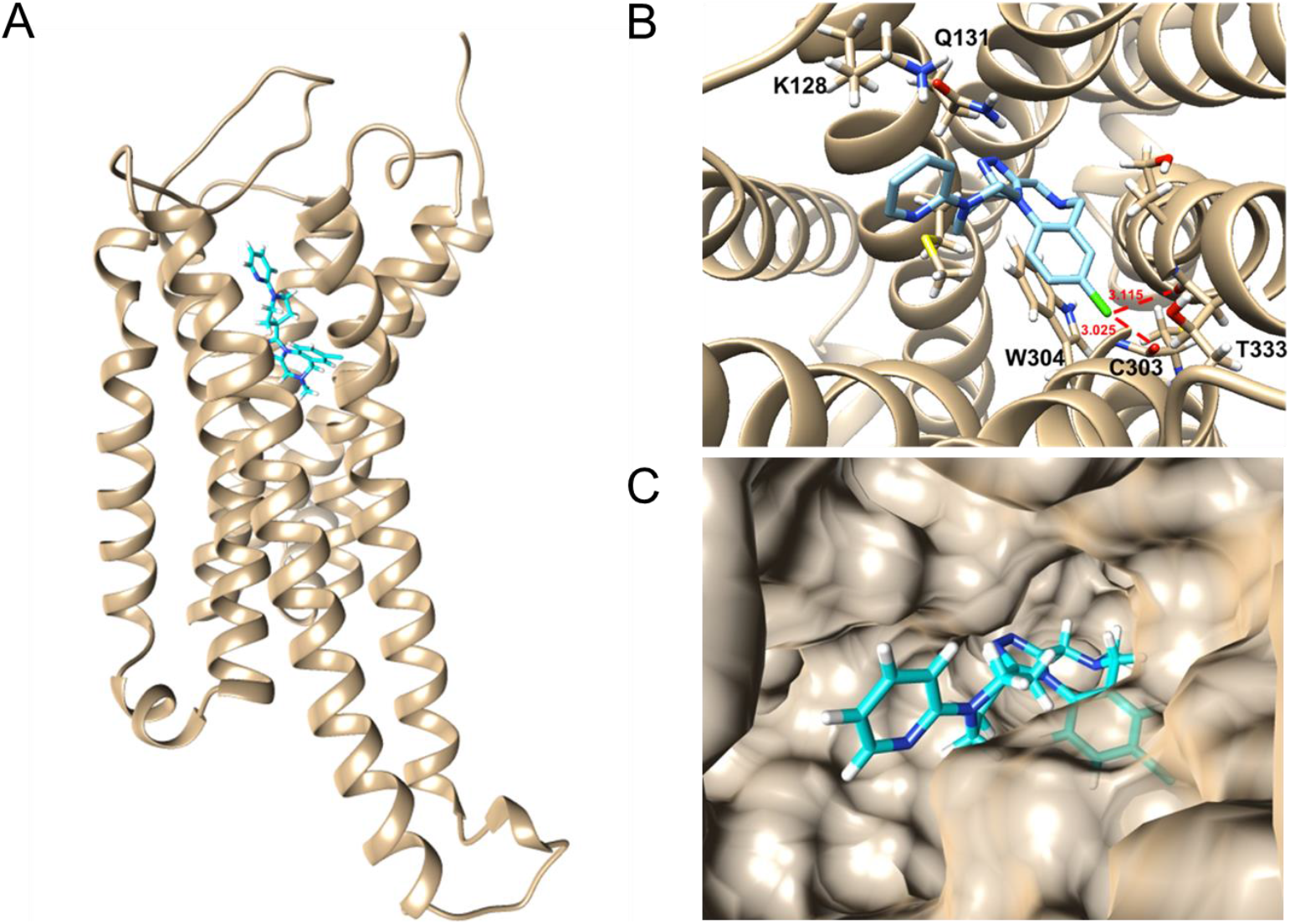
Molecular docking of PF-184563 to the V1A receptor. (**A**) Proposed binding site of the V1A receptor. (**B**) Predicted binding conformation of PF-184563 at the binding site. Residues within 3 Å are shown, with the distance of the chlorine atom with the carbonyls of C303 and T333 highlighted in red. (**C**) Surface representation of the binding pocket.

### Radiochemistry and Cell uptake assays

As depicted in **Figure 4A**, the amino group of precursor **19** was amenable for conventional carbon-11 labeling with [^11^C]CH_3_I, thus providing the radiolabeled analog, [^11^C]**17**, in excellent radiochemical purity (> 99%) and molar activities ranging from 37 - 46 GBq/µmol (n=5). [^11^C]**17** was synthesized with an automated module sequence that constituted radiolabeling, purification and formulation, with an overall synthesis time of 45 min. As expected, [^11^C]**17** was stable in saline containing 10% EtOH, and no degradation was observed for up to 90 min. Employing the shake-flask method, a logD_7.4_ of 2.13 ± 0.01 (n=3) was determined.

**Figure 4.**
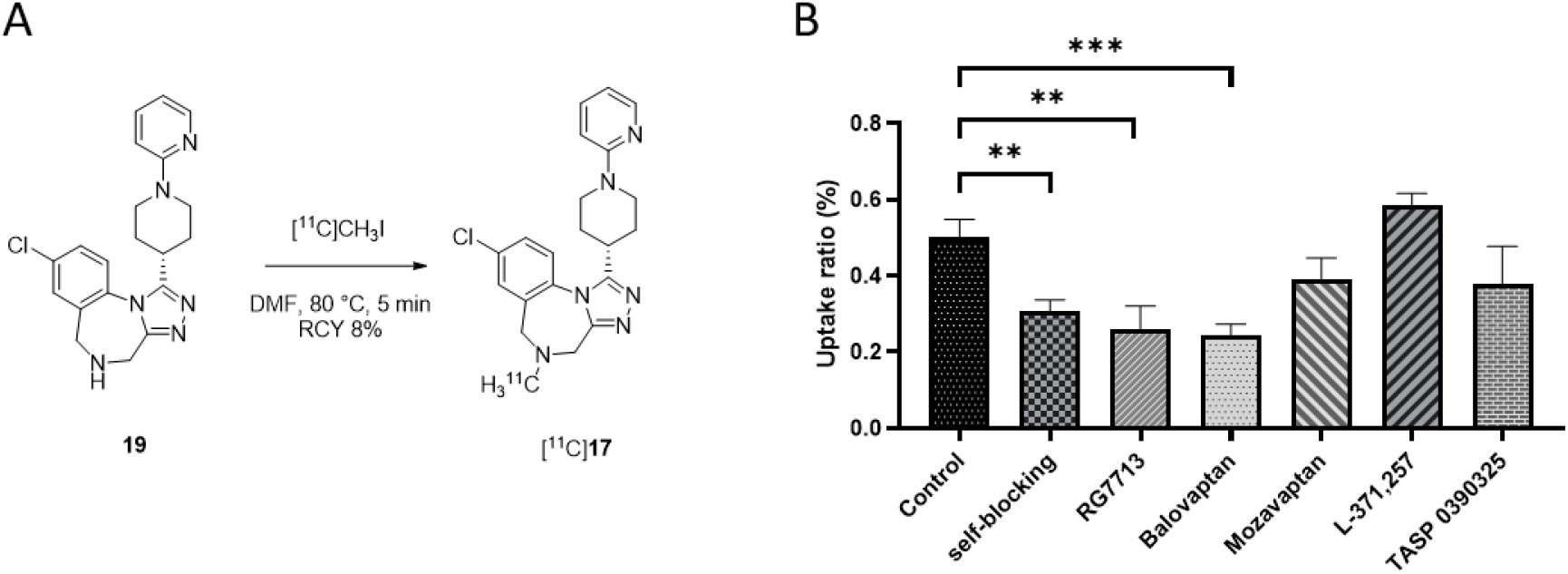
Radiolabeling and cell uptake studies with [^11^C]**17. A**. Radiosynthesis of [^11^C]**17**. RCY, non-decay corrected radiochemical yield. **B**. Uptake of [^11^C]**17** in a human V1A receptor Chinese hamster ovary (CHO) cell line. Uptake was significantly reduced in the presence of non-radioactive **17**, as well as the V1A receptor antagonists RG7713 and balovaptan. In contrast, no significant blocking effect was observed in the presence of mozavaptan (V2 receptor antagonist), L-371,257 (oxytocin receptor antagonist) and TASP 0390325 (V1B receptor antagonist). All blockers were used in a final assay concentration of 1 µM (>100-fold excess).

Cell uptake assays were conducted in the presence of different inhibitors to demonstrate the specificity and selectivity of [^11^C]**17** to the V1A receptor. As such, human V1A receptor-expressing Chinese hamster ovary (CHO) cells were seeded and incubated at 37 °C with either [^11^C]**17** (**Figure 4B**, control) or with a combination of [^11^C]**17** and a >100-fold excess of the non-radioactive blockers, **17** (self-blocking), V1A antagonists, RG7713 and balovaptan, the V2 antagonists, mozavaptan and L-371,257, as well as the V1B antagonist, TASP 0390325). A significant reduction of [^11^C]**17** cell uptake was observed under blockade conditions with all three V1A antagonists, while there was no significant blockade with mozavaptan, L-371,257 and TASP 0390325. These results indicate that [^11^C]**17** specifically binds to the V1A receptor *in vitro*, thereby exhibiting appropriate selectivity over V1B and V2 receptors.

### Autoradiography and PET experiments

Dynamic PET scans were acquired following tail vein injection of [^11^C]**17** in female CD-1 mice at the age of 8-10 weeks (n=4). Representative baseline sagittal and coronal PET images are depicted in **Figure 5A** and **5B**, respectively. In agreement with previous reports on high V1A receptor expression in the liver, we observed a high tracer uptake in this particular organ. To assess specific binding of the tracer to V1A receptors, commercially available V1A antagonist, balovaptan (3 mg/kg), was administered intravenously five min prior to tracer injection (**Figure 5C-D**). The liver was well delineated, exhibiting an appropriate signal-to-back ratio and allowing quantitative PET (**Figure 5E**, time-activity curves in the mouse liver). Of note, the high tracer uptake in the liver was substantially reduced when balovaptan was co-administered. Similarly, a high degree of specific binding was observed on autoradiograms of the rat liver, as shown in **Figure 5F**. In agreement with our findings, previous reports show that V1A receptor expression is abundant in hepatocytes, from which the V1A was first cloned by Morel et al. in the early nineties [42]. Despite the evident blockade effect, we did not achieve a full blockade in the liver, which points towards the presence of a small fraction of unspecific radio-metabolites. Whether the latter observation will persist in higher species remains to be elucidated.

**Figure 5.**
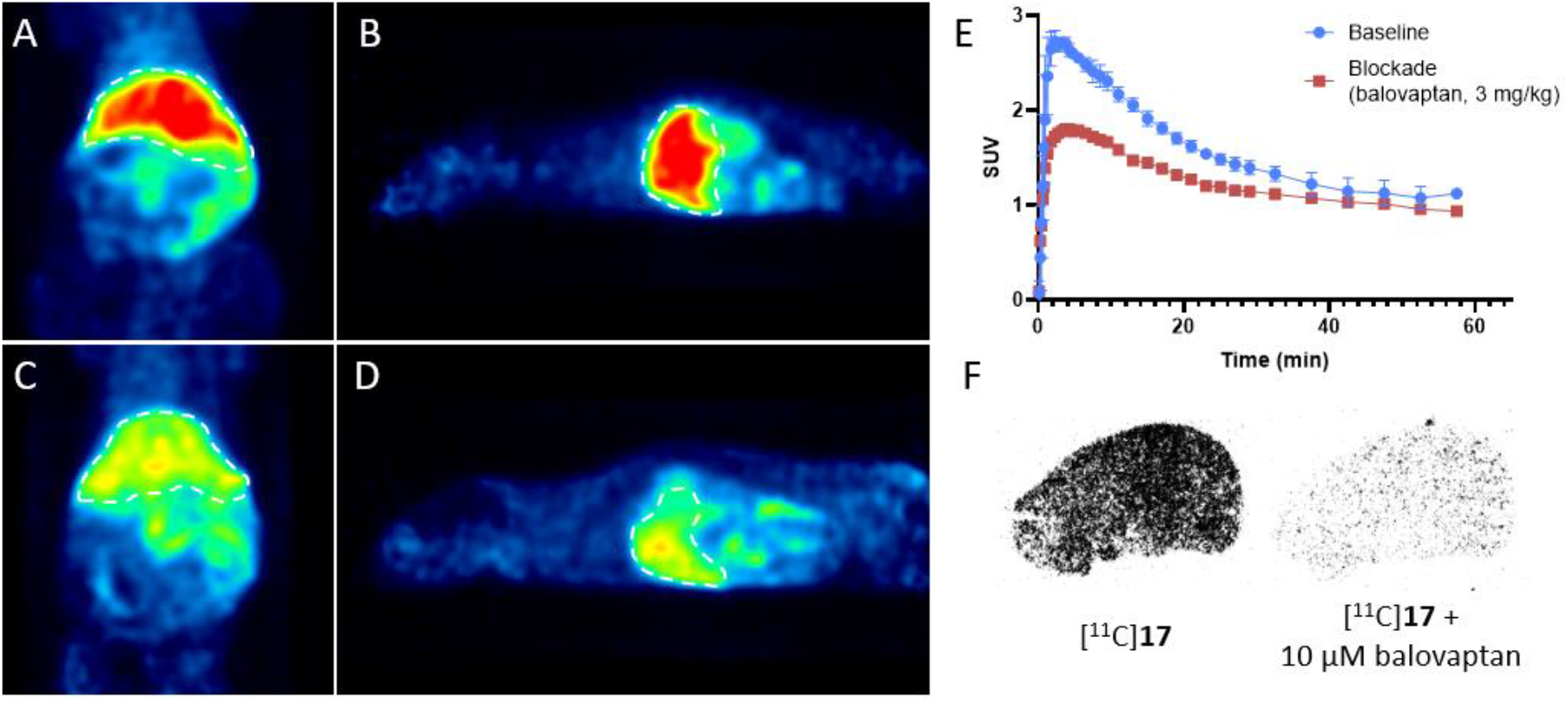
Rodent PET images and autoradiogram with [^11^C]**17**. Representative (**A**) coronal and (B) sagittal PET images of CD1 mice under baseline conditions (n=3), where only [^11^C]**17** was injected. Representative (**C**) coronal and (D) sagittal PET images under blocking conditions (n=1), where [^11^C]**17** and V1A antagonist, balovaptan (3mg/kg), were co-administered. Specific tracer binding to V1A was observed in the PET images (liver is highlighted by dashed white line) corroborated by the attenuated time-activity curve (**E**) in the liver under blockade conditions as well as by *in vitro* autoradiography (**F**) on Wistar rat liver sections. SUV, Standardized uptake value.

### Ex vivo biodistribution study

In a next step, we sought to quantify the tracer uptake in a comprehensive set of organs by *ex vivo* biodistribution. Biodistribution data of [^11^C]**17** was obtained in CD-1 mice at different time points (5, 15, 30 and 60 min) post injection. The data is presented as percentage of injected dose per gram tissue (**Figure 6**, %ID/g). While the radioactivity uptake was generally high in the liver, kidney and small intestine, we observed a sustained accumulation with relatively slow washout – particularly at >15 min post injection – in the pancreas, thyroid, spleen and the heart, indicating that there might be specific binding in those organs. Along this line, we performed a blocking study with the clinical V1A antagonist, balovaptan, to confirm the specificity of the signal. Indeed, a significant signal reduction under blockade conditions was corroborated for the spleen, heart and pancreas, thus suggesting that [^11^C]**17** is sensitive to V1A receptor expression in these organs *in vivo* (**Figure 7**). Compared to previously reported V1A PET ligands, [^11^C]**17** exhibited a relatively low brain uptake. [33] However, given that the objective was to assess peripheral V1A receptor expression, the low brain uptake did not hamper the successful outcome of the study. Low brain penetration of [^11^C]**17** may have been attributed to efflux by P-gp transporters located at the blood-brain barrier. Indeed, there have been reports linking the structural scaffold of **17** to P-gp transport. [28] In this study, efflux from the CNS may indeed have resulted in a beneficial effect on systemic availability of [^11^C]**17** and enhanced specific uptake in peripheral organs. Of note, our findings are in accordance with the known expression patterns of the V1A receptor across different peripheral organs. While Morel et. al demonstrated the presence of V1A receptors in the spleen and the kidney [42], Mohan et al. confirmed the expression of pancreatic V1A in murine islets, pancreatic rat cells (BRIN BD11) as well as human 1.1B4 beta-cells. [36] As for the cardiovascular system, the presence of V1A receptors was confirmed in cardiomyocytes [43] as well as in vascular smooth muscle cells, where it was shown to modulate blood pressure homeostasis and baroreflex sensitivity.[44]

**Figure 6.**
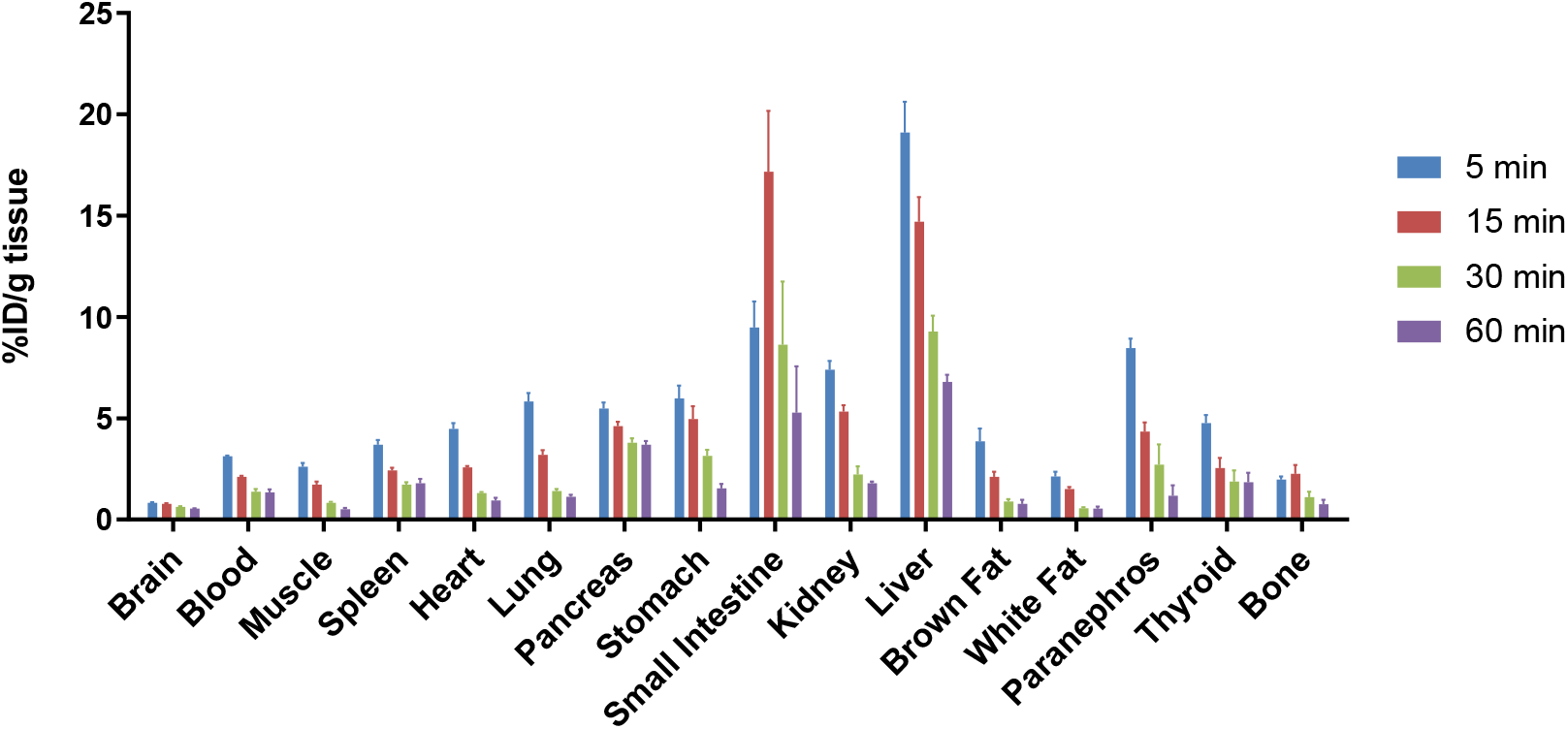
(A) *Ex vivo* whole body biodistribution in CD-1 mice at 5, 15, 30 and 60 min post injection of [^11^C]**17**. The results are expressed as the percentage of the injected dose per gram tissue (% ID/g); All data are presented as mean ± standard error (SE), *n* = 4.

**Figure 7.**
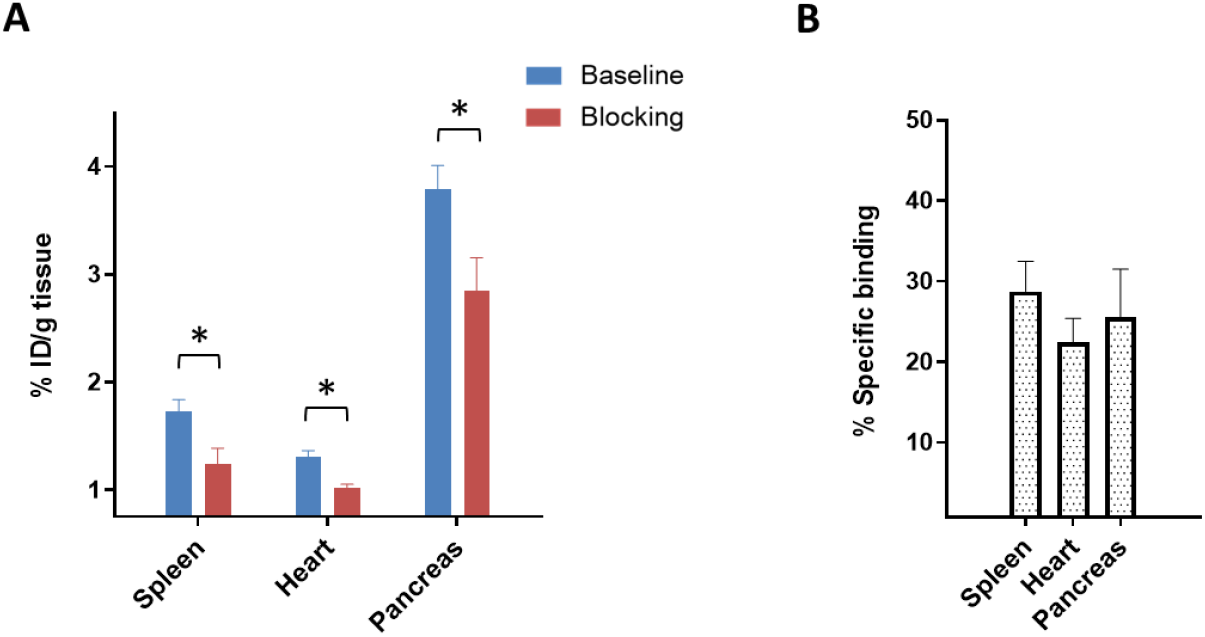
*Ex vivo* biodistribution in CD-1 mice under baseline and blockade conditions at 30 min post injection, with specificity in the spleen, heart and pancreas. Balovaptan (3 mg/kg) was used as a blocking agent. (**A**) Percentage of the injected dose per gram of wet tissue (% ID/g) under baseline and blocking conditions. A significant signal reduction was observed in the spleen, heart and pancreas. (**B**) Percentage of specific binding observed in each of the three organs. All data are mean ± standard error (SE), *n* = 4. Asterisks indicate statistical significance at *p* < 0.05.

A wealth of knowledge has been accumulated for the role of V1A receptors in peripheral pathologies such as liver cirrhosis, heart failure and diabetes. As such, V1A receptor agonists have been shown to promote liver regeneration in rats by triggering hepatic DNA and protein synthesis, ultimately resulting in an improved proliferation of healthy liver cells. [45] Further, the administration of dual V1A/V2 receptor antagonist, conivaptan, substantially attenuated water retention and dilutional hyponatremia in a rat model of liver cirrhosis. [46] Indeed, our newly developed PET probe may be harnessed (1) to assess target engagement of conivaptan on hepatic V1A receptors and (2) to monitor changes in V1A receptor density during pharmacological treatment, thereby ensuring sustained treatment efficacy as well as guiding potential changes in dose regimens.

V1A antagonists have further proved useful in cardiovascular disease (CVD). While the endogenous ligand, AVP, has been shown to be upregulated in patients with congestive heart failure, it is currently unclear whether V1A levels are altered in heart failure patients. The availability of a clinically validated V1A PET radioligand would allow studies to assess the utility of V1A as a disease biomarker and further support the development of suitable V1A antagonists for the treatment of heart failure in patients with elevated levels of AVP, given that selective cardiac V1A overexpression was shown to trigger cardiac hypertrophy and reversible left ventricular dysfunction in mice. [47]

V1A receptors have been implicated in diabetes mellitus. [48] Compelling evidence has been obtained from studies with V1A receptor knock-out mice that displayed significantly higher plasma glucose levels and exhibited a pre-diabetic phenotype with impaired glucose tolerance. [49] Along this line, blunted V1A expression levels in the pancreas or liver may serve as prognostic marker in pre-diabetic patients. If shown to predict disease progression and transition into diabetes, V1A targeted PET may indeed support contemporary diagnostic tools by further stratification into high-vs. low-risk patient populations.

## Conclusion

To improve our understanding on how V1A receptor expression and function is affected in peripheral organs such as the liver, heart, pancreas, spleen or kidney, a dedicated tool for non-invasive receptor visualization and quantification in patients is urgently needed. Notably, a clinically validated V1A-selective PET radioligand is currently lacking, and previous studies have exclusively focused on the development of CNS-targeted probes. To the best of our knowledge, this is the first report on a V1A PET radioligand that exhibits appropriate *in vitro* and *in vivo* specificity in peripheral organs, as evidenced by blockade studies in the liver, heart, pancreas and spleen. Our data suggests that [^11^C]**17** harbors potential to enable selective V1A-targeted imaging and warrants further studies in animal models of liver and cardiovascular diseases. The availability of a clinical V1A-selective PET radioligand would facilitate drug development via target engagement studies and contribute to an improved understanding of the versatile roles of V1A receptors, ultimately resulting in an improved diagnostic care for the patient.

## Experimental Section

Experimental procedures were generally performed as previously reported by our group, however, with minor modifications. [50, 51] All chemicals used for the synthesis of intermediates and target compounds were acquired from commercial vendors and then used without further purification steps. Silica gel was used for compound purification by column chromatography. Further, 0.25 mm silica gel plates were used for analytical thin layer chromatography (TLC). NMR spectra were obtained on a 300 MHz Bruker spectrometer, whereas “ppm” was used to indicate the chemical shifts (d) and “Hertz” was the unit of coupling constants. Multiplicities were assigned as follows: s (singlet), d (doublet), dd (doublet of doublets), t(triplet), q (quartet), m (multiple), or br (broad signal). Detailed chemical procedures and NMR spectra can be found in the supplemental information. Molar activities were assessed at the end of synthesis by comparison to a standard curve with known concentrations of target compound **17**. All animal studies were conducted in accordance with the guidelines of the Massachusetts General Hospital and were approved by the Institutional Animal Care and Use Committee. Animals were accommodated on a 12 h light/12 h dark cycle and were provided with food and water ad libitum.

### Molecular docking

The structure of **17** was generated with IQmol (IQmol, version 2.11), and minimized with the UFF force field. The structure of V1A was downloaded from the GPCR database (https://gpcrdb.org) which has 3 available homology models of the protein in the inactive, intermediate, and active state. The active form was used for initial molecular docking of **17** and prepared through the Dock Prep function of UCSF Chimera (UCSF Chimera, version 1.13.1), which involved the addition of hydrogen atoms, deletion of solvent, and assignment of AMBER ff14SB force field parameters for standard residues. 10 conformations were obtained from the molecular docking using AutoDock Vina 1.1.2, where the top binding conformation (best score) was reported. Energy minimization was performed using the Amber20 software. [39-41]

### Radiochemistry

Carbon-11 labeling was performed according to reported procedures. [52] Briefly, [^11^C]methyl iodide was synthesized from cyclotron-produced [^11^C]CO_2_, that was generated by the ^14^N(p, α)^11^C nuclear reaction. [^11^C]methyl iodide was transferred under heating with helium as a carrier gas into a reaction vial containing precursor **19** (1.0 mg, 2.6 µmol), and anhydrous DMF (200 μL). Radiosynthesis was performed at 80 °C for 5 min, followed by HPC purification on a Phenomenex Luna 5μ C18 column (10 mm i.d. × 250 mm) using a mobile phase of CH_3_CN / H_2_O (v/v, 35/65) containing 0.1% Et_3_N and at a flow rate of 5.0 mL/min. The retention time of [^11^C]**17** was 14.0 min. The radioactive fraction corresponding to the desired product was collected and passed through a C-18 light cartridge, eluted with 0.3 mL EtOH and the product was formulated with 5-10% EtOH in saline. The total synthesis time was ca. 45 min from end of bombardment (EOB). Radiochemical and chemical purity was measured by analytical HPLC (Xselect Hss T3 column, 4.6 mm i.d. × 150), whereas the identity of [^11^C]**17** was confirmed by the co-injection with non-radioactive **17**.

### Lipophilicity

LogD determination was conducted as previously reported. [51] Briefly, [^11^C]**17** was added to a mixture of n-octanol and aqueous PBS solution (0.1 M, pH 7.4). After vortexing for 5 min, samples were centrifuged for 5 min (3500 – 4000 rpm) to ensure proper phase separation. Subsequently, organic and aqueous layers were aliquoted, weighed, and the radioactivity was measured using a PerkinElmer 2480 Wizard automatic gamma counter. LogD was calculated based on the Log of the ratio between radioactivity in the n-octanol and PBS layer.

### Cell uptake studies

Human V1A Receptor-CHO cell line (CHO -V1a) was cultured in Ham’s F12 (Life Technologies) supplemented with 10% fetal bovine serum (Life Technologies) and Geneticin (G418,0.4 mg/mL, Life Technologies) with high humidity at 37°C and 5% CO2 for cell culture. Cells were subpassaged in a 1:2 split using 0.25% trypsin/0.02% ethylenediaminetetraacetic acid (EDTA). For cell uptake measurement, CHO -V1A cells were seeded into a 24-well plate at a density of 2×105 cells per well and incubated with 74 kBq/well of [^11^C]**17** at 37°C for 15, 30 and 45 min. Cells were then washed twice with cold PBS and harvested by adding 200 μL of 1 N NaOH. The blocking assay was preformed similarly to the procedure described above except that the non-radioactive reference compound, Rg7713, balovaptan, mozavaptan, L371257 or TASP0390325 were added into the wells and incubated for 60 min at 37°C before adding [^11^C]**17**. Cell suspensions were collected and measured in a gamma counter, whereas cell uptake, internalization and efflux were expressed as the percentage of the added dose (%AD) after decay correction. The data points of cell uptake and blocking studies constitute averages of quadruplicate wells.

### *In vitro* autoradiography

Autoradiography experiments were conducted as previously reported [53]. Briefly, liver tissue from male Wistar rats (8 weeks of age, n=2) was embedded in Tissue-Tek® (O.C.T.) and sections of 10 µm thickness were cut and stored at -20 °C until further use. Prior to the autoradiography experiments, liver sections were initially thawed for 10 min on ice and subsequently preconditioned for 10 min at 0 °C in assay buffer 1 (pH 7.4) containing 50 mM TRIS, 5 mM MgCl2, 2.5 mM EDTA and 1% fatty acid free bovine serum albumin. Upon drying, the tissue sections were incubated with [11C]17 for 15 minutes at 21 °C in a humidified chamber. For blockade, 10 µM of the clinically validated V1A receptor antagonist, balovaptan, was used. Liver slices were washed for 5 min in assay buffer 1 and further washed twice 3 min in assay buffer 2 (assay buffer 1 without bovine serum albumin). Tissue sections were dipped twice in distilled water and subsequently dried and exposed to a phosphor imager plate.

### PET imaging

PET imaging studies were conducted as previously reported. [54] Briefly, a G4 PET scanner (Sofie) was used to acquire PET scans, which were performed under 1%–2% isoflurane/air (v/v) anesthesia. Female CD-1 mice (8-10 weeks of age, n=4) were administered 1.5-1.9 MBq of [^11^C]**17** (dissolved in saline containing 5% ethanol and 5% tween 80, 100 µL per mouse) by use of a preinstalled tail-vein catheter. Dynamic PET images were acquired for 60 min in a 3D list mode. For blocking experiments, intravenous injection of the clinically validated V1A-selective ligand, balovaptan (3 mg/kg) dissolved in saline containing 5% ethanol and 5% tween 80 (100 µL), was performed 5 min before tracer injection. As previously described, ASIPro VW software was used for the reconstruction of the dynamic PET images, which are presented as average of the frames from 2-10 min post injection. Time-activity curves are presented as standardized uptake values, as previously reported [51].

### *Ex vivo* biodistribution

Biodistribution experiments in this work were conducted according to slightly modified previous reports. [51, 54] In brief, [^11^C]**17** (0.56 MBq/100 µL) was intravenously injected via tail-vein of female CD-1 mice (8-10 weeks of age, n=16). Animals were sacrificed by cervical dislocation at different time points (5, 15, 30 and 60 min) post [^11^C]**17** injection, whereas organs of interest were collected and weighted. We used a Packard Cobra 5002 automated Gamma Counter to determine the radioactivity in each organ. All experiments were conducted with four replicates and the data is presented as average ± standard error (SE) percent injected dose per gram wet tissue (%ID/g).

### Statistical analysis

Statistical analysis was conducted using either an unpaired two-tailed Student’s t test for comparisons of two independent values or a 1-way ANOVA for multiple comparisons as appropriate. Asterisks were used to indicate statistical significance: *P < 0.05, **P ≤ 0.01 and ***P ≤ 0.001.

## Compliance with ethical standards

### Funding

AH was supported by the Swiss National Science Foundation. LJY’s contribution was supported by NIH grants P50MH100023 to LJY and P51OD11132 to YNPRC.

### Author contributions

AH and SHL designed the research; AH, ZX, XX, JC, RSV, SK, CZ, JR, TEJ, TS, PR, AA and YS performed research; ZX and XX analyzed the data; AH wrote the paper; PR, AA, YS, CR, LJY, and SHL reviewed the paper.

